# Wounding promotes root regeneration through a cell wall integrity sensor, the receptor kinase FERONIA

**DOI:** 10.1101/2022.08.27.505519

**Authors:** Qijun Xie, Weijun Chen, Fan Xu, Shiling Ouyang, Jia Chen, Xuening Wang, Yirong Wang, Longfer Mao, Wenkun Zhou, Feng Yu

## Abstract

Wounding caused by various stresses is the initial event of plant regeneration. However, the mechanisms underlying the early wounding responses to promote plant regeneration remain largely unknown. Here, we report that the receptor kinase FERONIA (FER) interacts with Topless/Topless-related proteins (TPL/TPRs) to regulate the expression of regeneration-related genes to modulate root tip regeneration. One ligand of FER, rapid alkalinization factor 33 (RALF33), is stimulated by wounding and functions together with FER to promote regeneration. Single-cell sequencing data showed that the low-differentiation cell types in the stele may account for the enhanced regeneration ability in the *fer* mutant, especially in the columella and quiescent center (QC). Further interaction assays and analysis of the gene expression patterns in low-differentiation cell types confirmed that FER interacts with TPL/TPRs to regulate the expression of downstream regeneration-related genes. One of their downstream targets, an essential transcription factor (TF) in root regeneration, *ERF115*, acts downstream of FER-TPL/TPRs to control regeneration. Our results suggested a signaling pathway between the early wounding response and regeneration processes in roots.

**One-sentence summary:** RALF33-FER serves as an early signaling module between wounding and regeneration by functioning with TPL/TPRs in roots.

## INTRODUCTION

The bodies of both plants and animals are capable of repairing wounded tissues or organs, and this ability relies on a process termed regeneration. In plants, most organs have pluripotent cells that allow them to regenerate after wounding (1). After wounding, plants produce a series of second messengers (e.g., reactive oxygen species (ROS) and calcium) and elevate their levels of wound-related hormones (such as jasmonic acid (JA) and ethylene) (2, 3). Then, transcriptomic remodeling in cells around the wounding site is triggered, followed by the dedifferentiation and redifferentiation of these cells (4). A key transcription factor (TF) in the JA signaling pathway, MYC2, was reported to be an essential TF during wounding-induced regeneration, acting by binding directly to the promoter of two AP2/ERF TFs, namely, *ERF109* and *ERF115* (5). ERF109 also upregulates *ANTHRANILATE SYNTHASE* α1 (ASA1), a tryptophan biosynthesis gene in the auxin biosynthesis pathway, which may be a hormonal basis of regeneration (6–8). In addition, factors participating in regeneration processes, such as WINDs, PLTs, WOXs, and BOP1, have been identified (9–12). However, compared to the comprehensive identification of early wounding responses/signals (JA, ROS, etc.) and elements that control regeneration (ERF115, ASA, PLTs, etc.), the molecular basis or details of the signaling pathways that function between the early wounding responses/signals and regeneration processes remain largely unknown.

Receptor-like kinases (RLKs) are well known for their ability to transduce signals across membranes (13, 14). FERONIA (FER), a well-studied RLK, is known to respond to numerous stresses, including salt (15), temperature fluctuations (7), mechanical stress (16), and pathogens (17–19), reflecting its versatility in response to environmental cues. Importantly, FER is known to sense cell wall integrity during salt stress. The extracellular malectin-like domain of FER interacts directly with pectin in the cell wall. Salt stress-induced degradation of pectin was shown to be sensed by FER to trigger cell wall repair processes (15). Moreover, FER is functional in the development of several organs, such as leaves, cotyledons and seeds (20, 21). *FER* regulates seed size by inhibiting cell division during embryonic development (21). These observations suggest a potential role of FER in modulating cell differentiation after sensing wounding.

Herein, we report the participation of FER as a negative regulator in the early wounding signaling cascade to suppress root tip regeneration. One ligand of FER, rapid alkalinization factor 33 (RALF33), was found to accumulate in response to wounding and subsequently promote root regeneration. FER exhibited physiological interactions with Topless/Topless-related proteins (TPL/TPRs) to regulate the expression of *ERF115*. Based on our results, we present a model of the signaling cascade that occurs between early wounding signaling and root tip regeneration.

## RESULTS

### FER represses root tip regeneration

To determine whether FER is functional during wounding-induced regeneration, we performed root tip resection (22) and evaluated the regeneration rates of *fer-4*, a loss-of-function mutant of *FER* (23). Root tip resection allowed the removal of the root stem cell niche and meristem (Fig. 1A). The remaining stumps could regenerate the same organization based on the competence of the stump cells. After resection, *fer-4* presented a greatly enhanced capacity for root tip regeneration, especially type III resection, involving the removal of approximately 3/4 of the meristem (Fig. 1A-C). When the meristem was completely removed during type IV resection, the wild type (WT, Col-0) lost its regeneration capacity completely. Meanwhile, *fer-4* exhibited a considerable frequency of regeneration (Fig. 1 B-C). These data indicate that the *fer-4* roots had higher regeneration rates, which is surprising considering the weak features of the aboveground tissues of *fer-4* (e.g., small rosettes). The same trend was exhibited by *srn*, another mutant line of *FER* (C24 background) (23, 24). These results indicate that *FER* negatively regulates root tip regeneration.

**Fig. 1.**
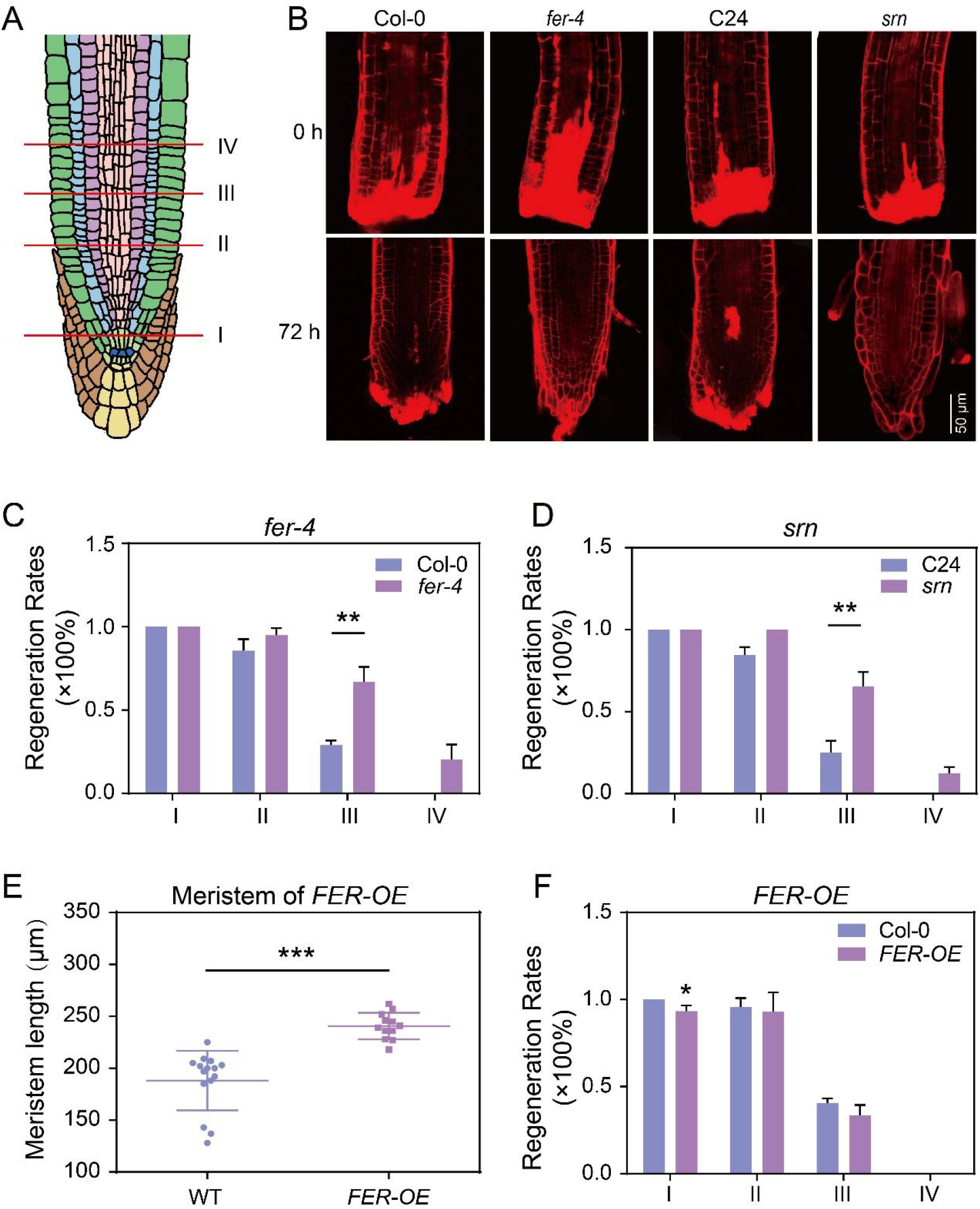
FER suppresses root regeneration. (A) Schematic diagram of the four types of resection. Type I refers to removal of root tips below (and including) the QC. Type II involves cutting off half of the meristem. Type III involves cutting off 3/4 of the meristem. Type IV refers to the removal of the whole meristem. (B) Representative root tips from Col-0, *fer-4*, C24 and *srn* at different time points after type III resection. Seedlings were stained with PI at 0 h and at 72 h after resection and were imaged by confocal microscopy. Bar = 50 μm. (C-D) Root tip regeneration frequencies 72 h after resection, described in (B). Bars represent the mean ± SE of 3 independent experiments, with at least 15 technical replicates per trial (Student’s t test, \*\**p* < 0.01). (E) Meristem length of the WT and *FER-OE* plants. The scatter dots shown indicate the meristem length of each independent root (Student’s t test, \*\*\**p* < 0.001). (F) Regeneration frequencies of Col-0 and *FER-OE* 72 h after resection. Bars represent the mean ± SE of 3 independent experiments, with at least 15 technical replicates per trial (Student’s t test, \**p* < 0.05).

We observed that *fer-4* and *srn* had enlarged meristems and increased columella cell layers (Fig. S1A-C). In parallel, EDU (5-ethynyl-2’-deoxyuridine) staining also revealed stronger division activity in the root tip of *fer-4* (Fig. S1D). Cell division also occurred widely in the elongation zone of *fer-4* (Fig. S1D). These findings led us to ask whether the stronger regeneration ability of *fer-4* resulted from the lower differentiation state. Correspondingly, *fer-4* exhibited many more lateral roots than the WT (Fig. S1E-F). It is likely that the lateral root-initiating cells of *fer-4* had stronger stemness or pluripotency, and the enhanced regeneration ability of *fer-4* may have been due to its lower differentiation state. However, overexpression of *FER* also led to increased meristematic size, yet the regeneration ability was slightly weaker than that of the WT (Fig. 1E-F). Therefore, the lower differentiation state of *fer-4* could not fully explain the elevation in regeneration competence. Otherwise, the enlarged meristem size in *FER-OE* should have increased the regeneration frequency. Overall, *FER* suppressed regeneration in roots after wounding.

### RALF33 responds to wounding to regulate root regeneration

A previous study indicated that injury caused by laser ablation did not affect the expression pattern of *FER* (3). Our data also showed that neither the expression pattern nor the protein abundance of FER was affected by resection (Fig S2A-B). On the other hand, the ligands of FER (i.e., RALFs) were reported to respond actively to environmental stimuli, and then to active or suppress FER (25). Interestingly, the root tip regeneration capacity of *llg1-2* (the loss-of-function mutant defective in a coreceptor of FER, Lorelei (lre)-like glycosylphosphatidylinositol (GPI)-anchored proteins (LLGs) (26) was clearly increased (Fig. S2C-D). These results imply that the molecular combination of RALFs and FER is required for root tip regeneration. To further determine whether a RALF (and if so, which RALF) is responsive to wounding and can regulate regeneration, we screened the *RALFs* that were expressed in the root meristem in the TAIR database. Accordingly, we obtained *RALF22, RALF23, RALF27, RALF31* and *RALF33* for further investigation. We also validated the expression pattern of the abovementioned RALFs by constructing GFP-tagged RALFs driven by their native promoters (Fig. S2E). Among them, *RALF33* showed high expression in most cell types in the root meristem, including the stem cell niche, lateral root cap, epidermis, cortex, endodermis, pericycle and steles (Fig. S2E). We then analyzed the root tip regeneration rates of the *RALF* overexpression lines (*RALF-OEs*). The *RALF22* overexpression line (*RALF22-OE*) exhibited slightly enhanced regeneration ability compared to the WT (Fig. 2A). In contrast to the subtle phenotype of *RALF22-OE, RALF33-OE* indeed showed a significant increase in its regeneration capacity (Fig. 2A-B). Since RALF33 was proven to be the ligand of FER (27), we hypothesized that the peptide RALF33 functions together with its receptor FER to regulate root tip regeneration. To verify this hypothesis, we exogenously applied 200 nM RALF33 to the stumps of the WT and *fer-4* after resection. The seedlings treated with RALF33 showed an increased regeneration rate compared to that of the control seedlings, yet RALF33 failed to promote the regeneration of *fer-4* (Fig. S3A-B). According to the expression profiles of *WOX5*, a marker of the quiescent center (QC), the mock seedlings exhibited a relatively broad expression pattern, indicating that the QC was not well formed (Fig. S3A). Meanwhile, in roots treated with RALF33, the narrow expression profile of *WOX5* was very similar to the pattern observed in intact roots, which showed 2 to 3 cells with pronounced *WOX5* expression. Correspondingly, *WOX5* in *fer-4* showed a narrow expression profile regardless of RALF33 treatment (Fig. S3A). The expression pattern of *WOX5* indicated that RALF33 accelerated QC regeneration. Interestingly, we observed many granular structures in the renewed columella cells of Col-0 (treated with 200 nM RALF33) and *fer-4* (with or without RALF33), with fewer structures observed in the Col-0 mock seedlings (Fig. S3A, indicated by red arrow). The granular structures observed were actually the starch granules in the columella, which are responsible for gravitropism. To confirm whether the enrichment of the granular structures observed reflected the improved renewal of the root caps, we performed a gravitropic response test 1 day after type III resection (22). Correspondingly, almost all the *fer-4* roots showed clear gravitropic bending 4 h after seedling rotation (Fig. S3C). Only ~16.2% of Col-0 seedlings showed a clear gravitropic response, but RALF33 treatment significantly increased this frequency to 41.3% (Fig. S3C). Therefore, the regeneration of columella cells in *fer-4* was significantly faster than that in Col-0, and RALF33 also promoted this process. In conclusion, RALF33 binds to its receptor FER to promote root tip regeneration, especially the QC and columella cells. Notably, treatment with RALF33 had a transient promoting effect on regeneration processes that was independent of the developmental stage.

**Fig. 2.**
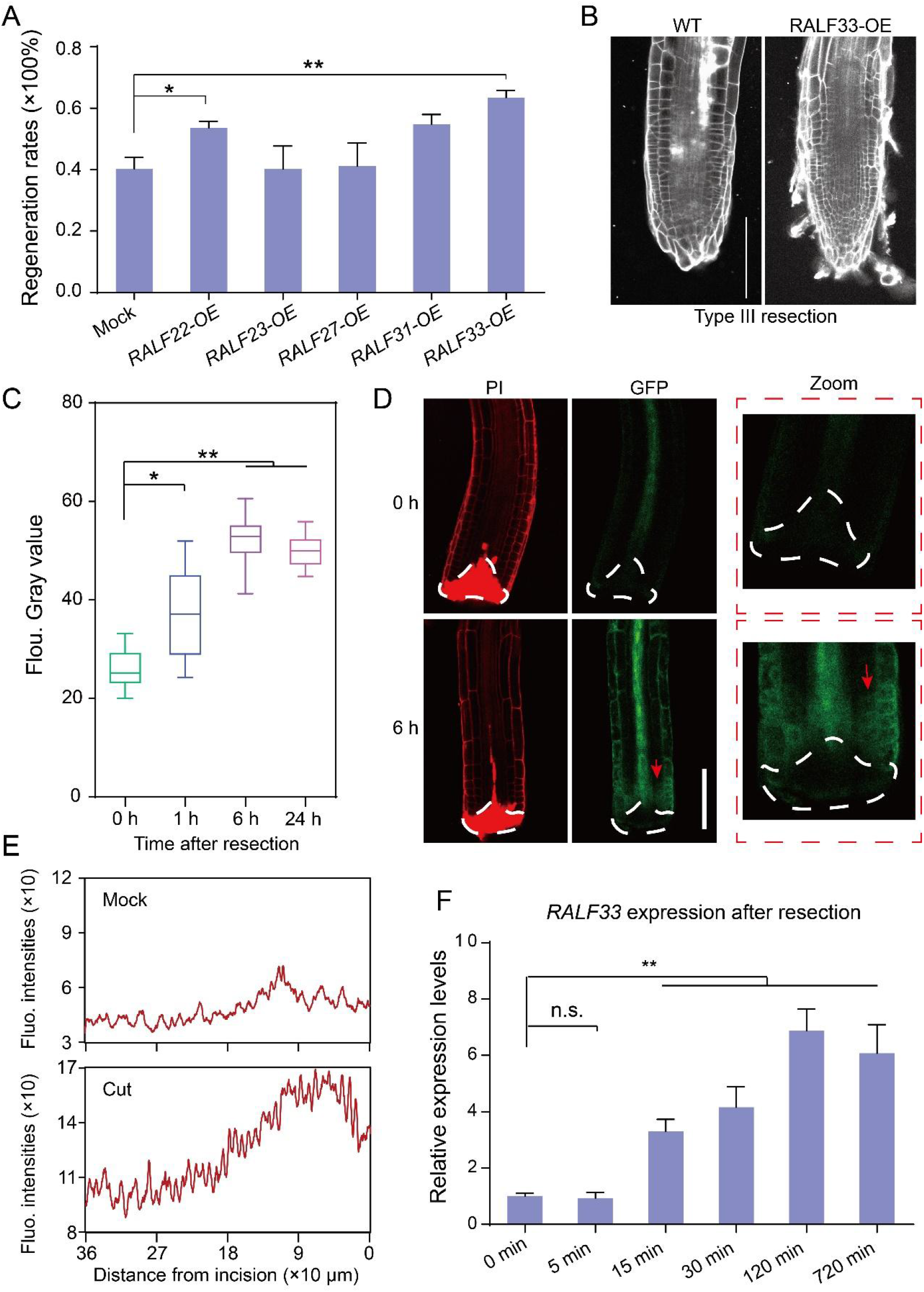
Wounding induces RALF33 accumulation to regulate regeneration. (A) Regeneration rates of the *RALF* overexpression lines following type III resection. Bars represent the mean ± SE of 3 independent experiments, with at least 15 technical replicates per genotype (one-way ANOVA, \**p* < 0.05; \*\**p* < 0.01). (B) Representative images of *RALF33-OE* and WT roots 3 days after type III resection. (C) RALF33 abundances at different time points after type III resection. Boxplot centers show median (n > 10 roots), and the box represents the interquartile range (one-way ANOVA, \**p* < 0.05; \*\**p* < 0.01). (D) Confocal images of *RALF33-GFP* seedlings following type III resection. The white dashed lines represent the cutting sites. The red arrow indicates the accumulated RALF33 near the wounding site. Bar = 100 μm. (E) Line profiles of GFP intensities at different distances from the incision. The line charts were generated from the images in D. (F) *RALF33* expression in roots at different time points after cutting was assessed by qRT–PCR. Each bar data point equals the mean ± SE of 3 independent experiments, with 3 technical replicates for each time point (one-way ANOVA, n.s., not significant; \*\**p* < 0.01).

We subsequently traced the dynamics of RALF abundance after wounding to determine whether RALF33 was responsive to wounding. Using the RALF-GFP marker lines, we found that RALF33 was more responsive to wounding than other RALFs (Fig. S4A-B). The RALF33 protein level increased rapidly (1 h) after resection and peaked at approximately 6 h (Fig. 2C-D). Importantly, we also observed RALF33 accumulation near the cutting site (Fig. 2D-E). The specificity of accumulation suggested an important role of RALF33 in the response to wounding.

Lags caused by de novo synthesis and GFP maturation have prevented the use of this marker line to study rapid dynamics that occur within a few minutes. To determine whether the upregulation of RALF33 was initiated quickly after resection, we quantified the expression levels of RALF33 at different time points via quantitative real-time PCR (qRT–PCR) (Fig. 2F). The increase in RALF33 expression was detectable as early as 15 min postresection and continued to increase in the subsequent 2 h. However, we did not detect changes within the first 5 min, when the early wounding signals (such as ROS) were active. The timing of RALF33 dynamics suggested that RALF33 may be located close to and relay the stereotypical signals of early wounding.

Collectively, we found that wounding induced rapid elevation in RALF33 levels in cells abutting the wounding site in the root tip, and RALF33 bound to FER to promote root tip regeneration. Notably, although only RALF33 was found to be responsive to wounding, we cannot rule out the possibility that other RALFs might be wound responsive.

### *FER* may regulate QC and columella regeneration by shaping the transcriptome of low-differentiation cells in the stele

We observed strong expression of a RALF (i.e., RALF33) in the stele (Fig. S2E). Interestingly, FER was also highly expressed in the stele (Fig S2A). These results led us to speculate about the important roles of RALF33 and FER in the stele. Importantly, we proved in a previous section that *RALF33-FER* could regulate the regeneration of QC and columella cells (Fig. S3). The regenerated columella cells originate from the stele (4); therefore, the physiology of stele cells is very likely shaped by *RALF33* and *FER*. To verify this hypothesis, we performed single-cell sequencing of the roots of the WT and *fer-4* (we will publish the detailed data from single-cell sequencing of *fer-4* in a separate work). The stele cells were classified into 4 clusters: pericycle, xylem, phenom and the low-differentiation type (which actually include the cambium, protoxylem and protophloem). Using the Spearman correlation coefficient of the transcriptome as an indicator, we discovered that the transcriptomes of cells with low differentiation in *fer-4* were greatly different from those in the WT-the correlation coefficient of the low-differentiation cell type in the above two genotypes was only 0.45 (Fig. 3A) and was at the same level as those between different cell types (e.g., the xylem and phloem in the WT). Consistent with the results revealed by correlation analysis, the number of differentially expressed genes (DEGs) in low-differentiation cells (normalized to the total number of detected genes) were approximately 4 times greater than those in xylem and phloem (Fig. 3B). We applied GO enrichment of the DEGs in low-differentiation cells from *fer-4* and WT. Seven of the GO terms among the top 20 terms were related to either wounding or regeneration (Fig. 3C). Therefore, transcriptomic variation in low-differentiation cells very likely contributed to the different regeneration rates of Col-0 and *fer-4*, especially that in the columella and QC.

**Fig. 3.**
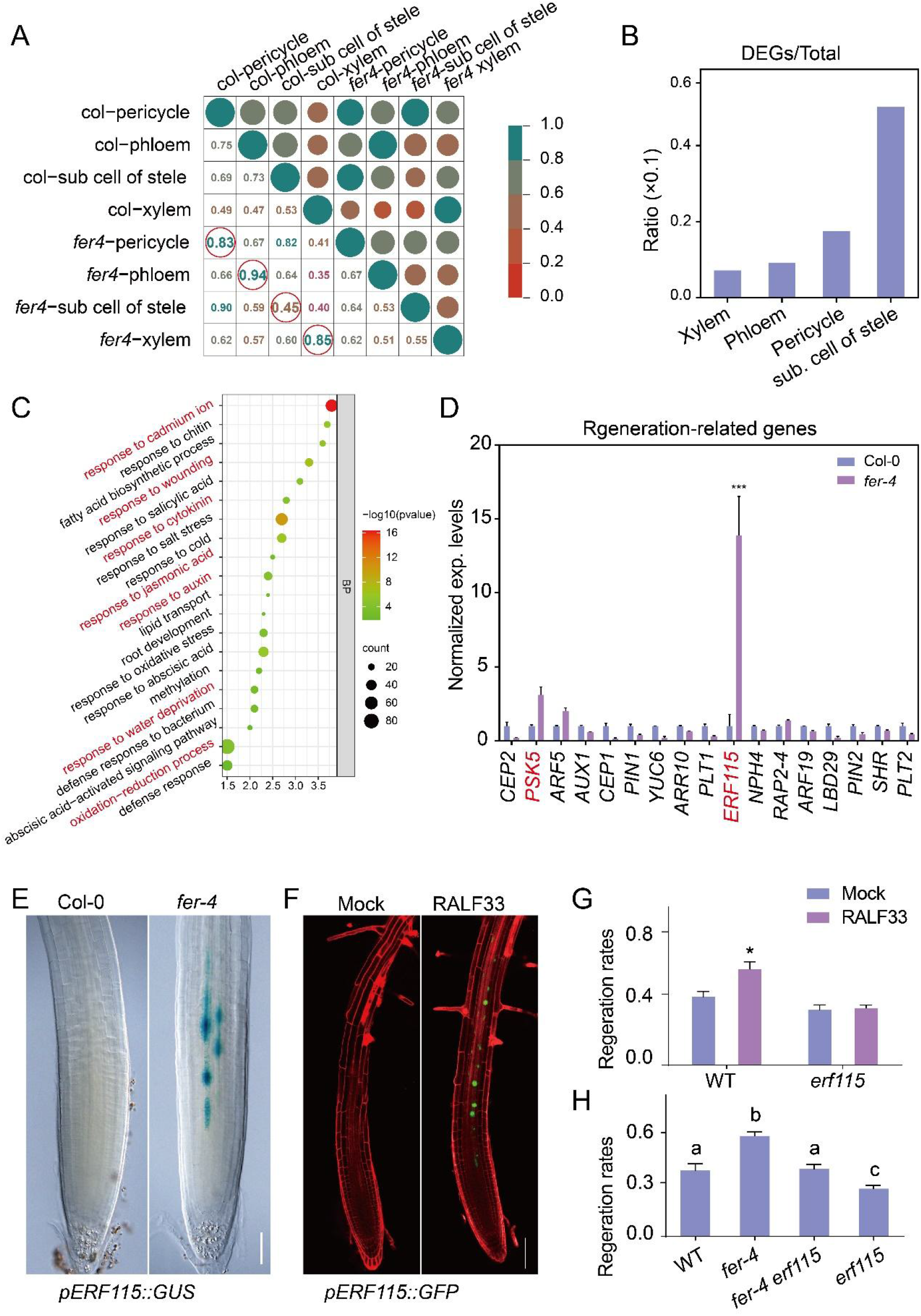
Single-cell RNA-seq analysis of cell clusters in the stele and ERF115 functional downstream of RALF33. (A) Spearman correlation coefficient of the transcriptome in different cell types from Col-0 and *fer-4*. Correlation coefficients are indicated by circle areas and colors of each in the top right quadrant and can be directly assessed by the numbers in the lower left quadrant. (B) DEGs/Total ratio in different cell types. ‘DEGs’ is the numbers of differentially expressed genes in Col-0 and *fer-4*. ‘Total’ is the number of total detectable genes. (C) Top 20 GO terms of DEGs in low-differentiation cells of Col-0 and *fer-4*. The GO terms marked in red are those related to wounding or regeneration. (D) Normalized gene expression levels of root regeneration-related genes in roots. Expression levels were obtained from RNA-seq data of Col-0 and *fer-4* roots (25), with 3 biological replicates. The expression level in *fer-4* was normalized to that in Col-0 (Student’s t test, \*\*\**p* < 0.001). (E) *ERF115::GUS* activities in roots of Col-0 and *fer-4*. (F) RALF33 induces the expression of *ERF115*. The images presented here are of the *pERF115::GFP-GUS* line following RALF33 treatment. (G) Regeneration rates of *erf115* in response to RALF33 treatments. RALF33 (200 nM) was applied to the stumps of Col-0 and *erf115* after type III resection. Bars represent the mean ± SE of 3 independent experiments, with at least 15 technical replicates per trial (Student’s t test, \**p* < 0.05). (H) Regeneration rates of the WT, *fer-4*, *fer-4 erf115* and *erf115* following type III resection. Bars represent the mean ± SE of 3 independent experiments, with at least 15 technical replicates per trial. The different lowercase letters indicate statistical significance (one-way ANOVA).

### RALF33-FER regulates root tip regeneration through ERF115

A previous review that summarized information on the genes currently known to be involved in root regeneration (28). We also examined the expression of these genes in the low-differentiation cells from Col-0 and *fer-4*. In total, 19 regeneration-related genes were differentially expressed (Fig. S5A). Among them, *ERF115* and its downstream peptide *PSK5* were upregulated (Fig. S5A). In addition, among the regeneration-related genes in our RNA-seq data for roots of the WT and *fer-4, ERF115* and *PSK5* were markedly enriched in *fer-4* (Fig. 3D). We confirmed this result using the *pERF115::GUS* reporter line in the Col-0 and *fer-4* backgrounds (Fig. 3E). The expression of *ERF115* was distinctly higher in the stele of *fer-4* (Fig. 3E). RALF33 treatment also stimulated *ERF115* expression, as revealed by using the *pERF115::GFP* marker line (Fig. 3F). ERF115 is an essential factor for root tip regeneration, and overexpression of *ERF115* significantly promotes root tip regeneration (5, 29). Therefore, we postulated that *ERF115* could function downstream of the RALF33-FER signaling module to regulate root tip regeneration.

To demonstrate whether *ERF115* acts downstream of RALF33-FER, we applied 200 nM RALF33 to the excised WT and *erf115* roots. Consistent with previous reports (5, 29), the regeneration frequency of the *erf115* mutant was significantly lower than that of the WT (Fig. 3G). Meanwhile, RALF33 treatment failed to elevate the regeneration frequency of *erf115* (Fig. 3G).

We also generated a *fer-4 erf115* double mutant to determine whether *ERF115* acts downstream of *FER*. Although the defective root hairs of *fer-4 erf115* resembled those of *fer-4* (Fig. S5B), the meristem size of the double mutant was reduced compared to that of *fer-4* (Fig. S5C-D). Hence, the enlarged meristem rather than the abnormal root hairs of *fer-4* could be attributed to the perturbation of *ERF115*. The regeneration rate of *fer-4 erf115* was also reduced to a level similar to that of the WT (Fig. 3H). In conclusion, ERF115 functions downstream of RALF33-FER to regulate root tip regeneration.

### TPL/TPRs serve as potential upstream regulators of ERF115

Transcriptional regulation by ERF115 of its downstream targets occurs in the nucleus, yet FER is located on the plasma membrane (30). Spatial compartmentalization prevents FER from directly regulating *ERF115*. FER most likely regulates *ERF115* through its interacting proteins. Based on the results of our previous yeast two-hybrid (Y2H) screening (7), we focused on the transcriptional corepressor TOPLESS-RELATED 1 (TPR1). TPR1 belongs to the TPL/TPR protein family. The first identified member, TOPLESS (TPL), is known as a master regulator of root fate determination (31). In addition, TPL/TPRs also play an important role in maintaining the stemness of columella stem cells (32). These reports emphasized the possible role of TPL/TPRs in regeneration, because cell fate regulation via either root fate determination or stemness maintenance indicates the cytological nature of regeneration. Interestingly, TPR1 distributes in plasma, cytoplasm and nuclear (Fig. S6A). The subcellular localization of TPR1 also makes it possible to mediate the transcription regulation of *ERF115* by FER.

As gene expression coregulators, TPL/TPRs have many downstream targets (33). To further confirm the possibility that FER regulates gene expression by interacting with TPL/TPRs, we performed TF enrichment analysis in the iGRN and PlantTFBD databases (see Materials and Methods). TPL/TPRs are actually not TFs but can modulate gene expression by recruiting other TFs. Hence, we alternatively focused on the enrichment of proteins interacting with TPL/TPRs. By searching iGRN (http://bioinformatics.psb.ugent.be/webtools/iGRN/) (34) and PlantTFBD (http://planttfdb.gao-lab.org/) (35) using 1507 regeneration-related DEGs in low-differentiation cell clusters Fig. 2C), we obtained 1260 and 567 TFs, respectively (Fig. S6B). The two shared 532 overlapping TFs, and 14 proteins among the 53 reported interacting proteins of TPL/TPRs were present among these 532 overlapping TFs. Therefore, certain DEGs in the low-differentiation cells from the WT and *fer-4* were likely regulated by TPL/TPRs. We also analyzed the transcriptomic overlap of *fer-4* (36) and *tpltpr1tpr4* (33). In total, 1908 overlapping genes among the DEGs of *tpltpr1tpr4* and *fer-4* were identified (Fig S6C). The high degree of overlap supported the hypothesis that FER interacts with TPL/TPRs to regulate the expression of its downstream genes. GO enrichment analysis using the 1908 overlapping genes revealed many GO terms related to wounding, such as response to wounding and response to jasmonic acid (Fig. S6D). The above bioinformatic analysis strongly suggested that FER regulates downstream genes (e.g., *ERF115*), especially those related to wounding, through TPL/TPRs.

### FER interacts with TPL/TPRs to promote their degradation

To verify the hypothesis from the previous section, we examined the interactions between FER and TPL/TPRs. We cloned the full-length CDSs of the five TPL/TPR family members, namely, TPL, TPR1, TPR2, TPR3 and TPR4 (TPL/TPR-AD), into an activation domain (AD)-containing vector. The TPL/TPR-AD constructs were cotransformed with FER-CD-BD, a recombinant vector with the cytosolic domain of FER (FER-CD) fused to the binding domain (BD), to perform a Y2H assay. The Y2H assay clearly revealed the interactions of TPR1, TPR3 and TPR4 with FER-CD, as well as a weak interaction of TPL (Fig. 4A). We further examined the interaction of TPR1 using a split-luciferase system. An interaction between TPR1 and FER-CD was demonstrated, verifying the interaction between FER and TPL/TPRs (Fig. 4B). We also performed a coimmunoprecipitation assay to study their interaction *in vivo*. The TPL/TPRs coimmunoprecipitated with FER-Flag (Fig. 4C). For the pull-down assay, we first immunoprecipitated the TPR1-Myc protein from Myc-tagged transgenic seedlings. The TPR1-Myc protein could be pulled down by FER-GST, which indicated that FER is physiologically associated with TPR1 (Fig. 4D). The *in vitro* phosphorylation assay also revealed that the kinase domain of FER was sufficient for phosphorylating the N-terminus of TPR1 (Fig. 4E). Overall, FER interacted with TPL/TPRs both *in vivo* and *in vitro*.

**Fig. 4.**
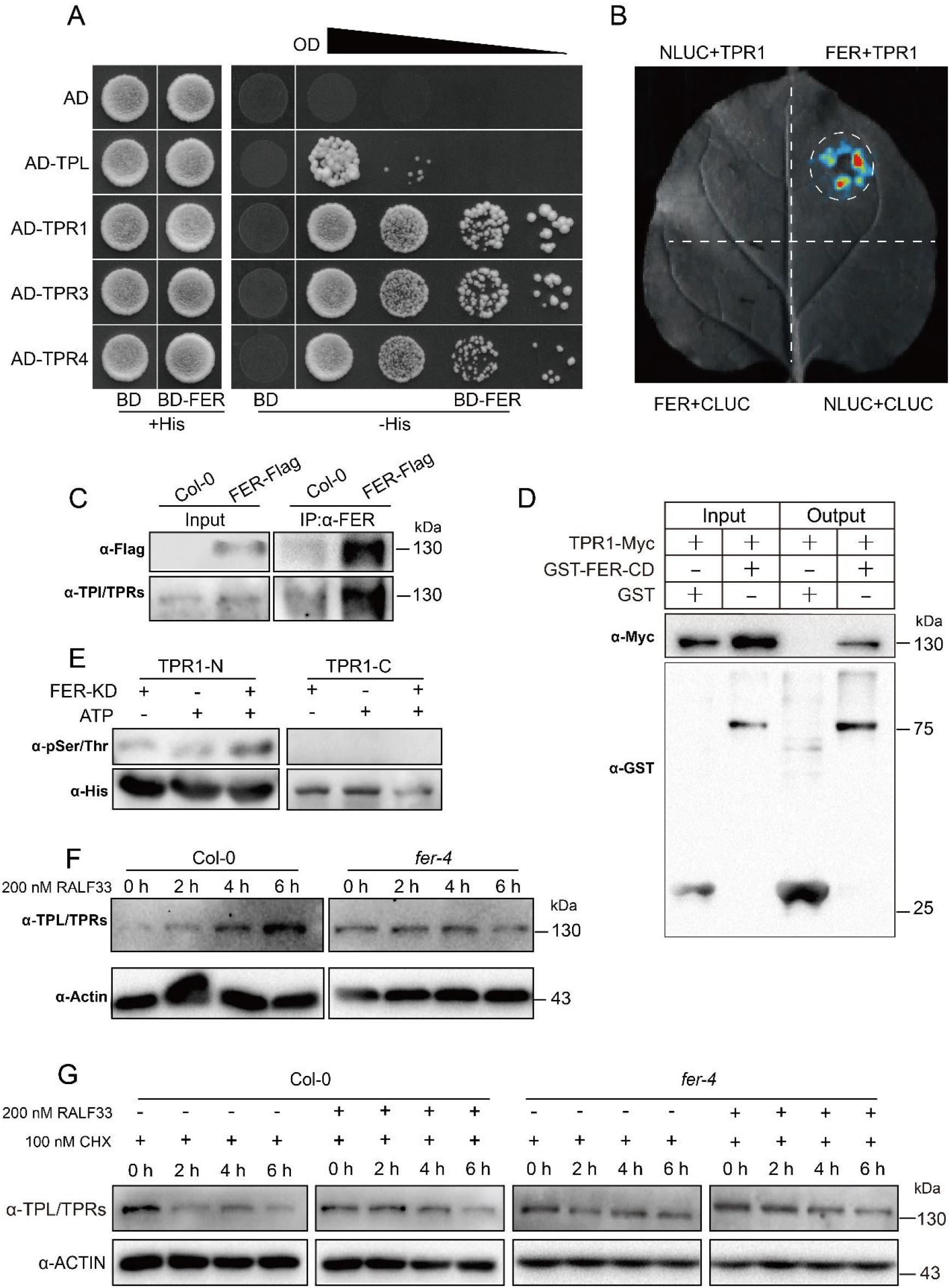
FER interacts with and phosphorylates TPL/TPRs. (A) Yeast two-hybrid assay of FER and TPL/TPRs. Synthetic dropout medium (-His) containing 20 mM 3-amino-1,2,4-triazole was used to examine the interaction. (B) Split-luciferase assay exhibiting the interaction between FER and TPR1. The cytosolic domain of FER and full-length TPR1 were used to test the interaction. (C) Co-IP assay showing the interaction between FER and TPL/TPRs. Protein lysates from FER-Flag and Col-0 seedlings were immunoprecipitated by anti-Flag magnetic beads, and interactions were detected by an TPL/TPR antibody. (D) GST pull-down assay of TPR1-myc and FER-CD. (E) In vitro phosphorylation assay of TPR1-N and TPR1-C by FER-KD. (F) TPL/TPRs abundances in Col-0 and *fer-4* at different time points following RALF33 treatments. (G) Stability of TPL/TPRs in Col-0 and *fer-4* after RALF33 treatment. Protein stabilities were assessed using CHX to block de novo protein synthesis.

What is the molecular significance of the interaction between FER and TPL/TPRs? To answer this question, we investigated the protein level of TPL/TPRs in response to RALF33 treatment in the WT and *fer-4*. We found that TPL/TPRs accumulated at significant levels in the *fer-4* mutant (Fig. 4F). Second, exogenous application of RALF33 led to the accumulation of TPL/TPRs in Col-0 but not in *fer-4* (Fig. 4F). Finally, when 100 μM proteasome inhibitor, cycloheximide (CHX), was applied to inhibit protein synthesis, the TPL/TPR levels were decreased more rapidly in the WT than in *fer-4* (Fig. 4G). Taken together, the results showed that FER interacts with TPL/TPRs to reduce their half-life.

### FER interacts with TPL/TPRs to regulate root tip regeneration

Does the increase in TPL/TPR abundance affect regeneration? To answer this question, we performed resection to investigate the regeneration of the triple mutant *tpltpr1tpr4* and performed overexpression of *TPR1 (TPR1-OE;* Fig. 5A). The *tpltpr1tpr4* mutant showed an attenuated regeneration frequency versus the WT, while *TPR1-OE* exhibited a stronger regeneration capacity (Fig. 5A-B). Application of 200 nM RALF33 to the resected *tpltpr1tpr4* stump demonstrated that TPL/TPRs function downstream of RALF33 to regulate regeneration, as *tpltpr1tpr4* was less sensitive to RALF33 (Fig. S7A-B). Hence, TPL/TPRs act as signal repeaters downstream of RALF33-FER to ultimately regulate regeneration. We also crossed *fer-4* with *tpltpr1tpr4* and obtained *tpr1fer-4* and *tpr1tpr4fer4*. the regeneration rates of *tpr1fer-4* and *tpr1tpr4fer-4* were lower than that of *fer-4* (Fig. 5C-D). Collectively, the results show that RALF33-FER interacts with Col-0 and *fer-4* to regulate root tip regeneration.

**Fig. 5.**
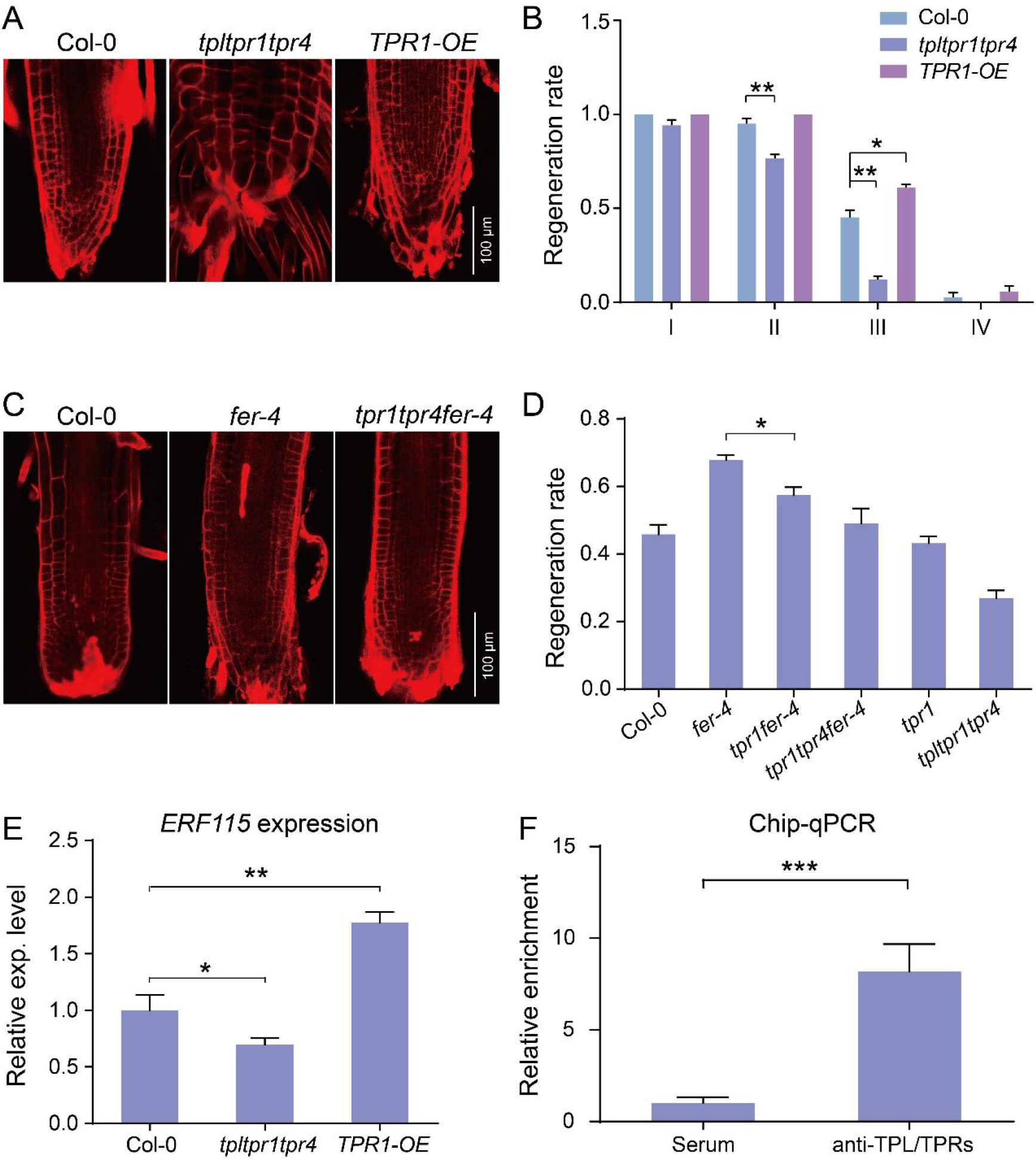
TPL/TPRs promote ERF115 expression to regulate root regeneration. (A) Representative images of Col-0, *tpltpr1tpr4* and *TPR1-OE* following 72 h of type III resection. (B) Regeneration rates of Col-0, *tpltpr1tpr4* and *TPR1-OE*. Bars represent the mean ± SE of 3 independent experiments, with at least 15 technical replicates per trial (one-way ANOVA, \**p* < 0.05; \*\**p* < 0.01). (C) Representative images of Col-0, *fer-4* and *tpr1tpr4fer-4* following 72 h of type III resection. (D) Regeneration rates of Col-0, *fer-4, tpr1fer-4, tpr1tpr4fer-4, tpr1*, and *tpltpr1tpr4*.Bars represent the mean ± SE of 3 independent experiments, with at least 15 technical replicates per trial (one-way ANOVA, *\*p* < 0.05). (E) *ERF115* expression levels in Col-0, *tpltpr1tpr4* and *TPR1-OE* cells. Expression levels in *tpltpr1tpr4* and *TPR1-OE* were normalized relative to that in Col-0. Bars represent the mean ± SE of 3 independent experiments, with 3 technical replicates per trial (one-way ANOVA, \**p* < 0.05; \*\**p* < 0.01). (F) ChIP analysis of the recruitment of TPR1 to the promoter of *ERF115*. ChIP was performed using an anti-TPL/TPRs antibody. DNA quantification using qRT–PCR. Bars represent the mean ± SE of 3 independent experiments, with 3 technical replicates per trial (Student’s t test, \*\*\**p* < 0.001).

### TPL/TPR regulates the expression of *ERF115*

We asked whether TPL/TPRs could regulate *ERF115* expression. Using qRT–PCR, we revealed a lower *ERF115* expression level in *tpltpr1tpr4* cells than in *TPR1-OE* cells (Fig. 5E). Moreover, TPL/TPRs interacted with the promoter of *ERF115* to regulate its expression, as indicated by ChIP–qPCR (Fig. 5F). This result is consistent with their corresponding phenotypes, which suggested that TPL/TPRs promote root tip regeneration (Fig. 5A-B).

## DISCUSSION

Based on our results, we proposed the following model: RALF33-FER can respond to wounding and transduce wounding-related signals by interacting with TPL/TPRs to ultimately regulate the expression of ERF115, a key TF involved in root tip regeneration (5, 29). The most significant result of this study is the elucidation of a molecular pathway serves as an early signaling module between wounding and regeneration.

Wounding-induced regeneration actually includes two processes: wounding and regeneration (37). However, to date, studies have mainly investigated these two events independently (28). Knowledge of the molecular basis of these two events is scarce. Our research shed light on RALF-FER as a functional module involved in this process. In our proposed model, the RALF33 protein responds within 1 h, and *RALF33* mRNA can respond to wounding in tens of minutes. Since the initiation of regeneration usually takes several hours (22, 28), it is quite reasonable that RALF33 serves as an earlier signaling molecule that transduces wounding-related signals for regeneration.

According to a previous publication, FER can sense wounding (15). What is the physiological significance of employing RALF33 to respond to wounding, since it seems achievable with FER alone? Importantly, cutting alone causes disruption of the cell wall in a single layer of cells near the incision, yet regeneration can be observed broadly around the wounding site (22). However, this phenomenon can be better explained if we take RALFs into consideration. Because RALFs are diffusible, they cause the regeneration of the surrounding cells by diffusing into intact cells that are distant from the incision (Murphy and De, 2014). Efforts should be made in the future to reveal the mechanism by which RALF33 and FER are regulated by wounding.

The regenerated root tip originates not from specific cryptic stem cells but multiple tissues from the remaining stump (4). Specifically, cells in the stele are respecified into stem cells to generate the QC and columella (4). It remains unknown whether this process is independent of the regeneration of other cell types (such as the cortex), and the mechanism underlying the regeneration of the QC and columella is unknown. The expression patterns of *RALFs* and *FER* suggested their important role in the stele (Fig. S2A, E). The accelerated QC and columella regeneration clearing reflected the differences in the physiology of stelar cells. We further revealed the difference in the distinct transcriptomes of low-differentiation cells in the stele. Therefore, it is likely that RALF-FER controls the regeneration of the QC and columella by shaping the transcriptome of low-differentiation cells in the stele. Our results provide a possible model to explain how a certain cell population is regenerated.

A previous publication indicated that TPL/TPRs act as corepressors (38). Here, we found that TPL/TPRs promoted the expression of *ERF115* and that the expression level of *ERF115* in *tpltpr1tpr4* was decreased. Unfortunately, we have not make further efforts to resolve this contradiction. It is not that surprising since there are many studies on the bilateral effects of transcription regulators (39). Take PIFs as an example; an individual gene may respond to PIFs inversely, and the same PIF may either up- or downregulate the expression of different genes (39).

In conclusion, the signaling pathway between wounding and regeneration still requires future investigation. Our research introduces the role of RLKs in this pathway. These processes also represent the checkpoints between wounding and downstream reactions, such as regeneration. Elucidation of this mechanism could provide molecular targets for genetic manipulation and improvement by, for example, grafting, cutting, and callus induction. We believe that further exploration of this topic will be helpful and yield promising results.

## MATERIALS AND METHODS

For full and detailed methods please se *SI Appendix, Materials and Methods*.

### Plant materials and growth conditions

The *Arabidopsis thaliana* ecotype C24 was used as a wild-type control for *srn*. In addition, Col-0 was the control for other mutants. The loss-of-function mutant *fer-4* was described previously (40). The *llg1-2* (*CS66106*) mutant was kindly provided by Doctor C. Li (Li et al, 2015). The *erf115* (*SALK_021981*) mutants, *ERF115-GUS* and *ERF115-GFP-GUS* were used in previous research (5, 29). The triple mutant *tpltpr1tpr4* and TPR1-OE were provided by Doctor J.B. Jin (41). *ERF115-GFP-GUS* (referred to as *ERF115-GFP* in the text) has been reported previously (5, 29).

To generate the GFP-tagged reporter line of *RALF22 (AT3G05490), RALF23 (AT3G16570) RALF27* (AT3G29780), *RALF31 (AT4G13950*), and *RALF33* (*AT4G15880*), DNA fragments of GFP-tagged full-length RALF CDSs under the driven by their native promoter were cloned into the pCAMBIA1300 backbone. The tagged *pCambia-1300-pRALFs::RALFs-GFP* constructs were then transformed into the Col-0 ecotype to generate the corresponding seedlings. The RALF overexpression constructs were generated by introducing the full-length CDSs carried by the pDT1 backbone into Col-0.

Arabidopsis seeds were surface sterilized by treating with 75% ethanol for 5 min followed by sodium hypochlorite for 15 min. The samples were washed 5 to 6 times with sterilized deionized water and sown on half-strength MS medium (1/2 MS medium) with 1% sucrose and 1% agar. The seeds were stratified in the dark at 4°C for 2 days and were subsequently transferred to a growth chamber under controlled conditions. The parameters of the growth chamber were set as follows: 22°C, 80% (relative humidity), 16/8-h light/dark.

### Root tip resection

The root tip resection method was based on the description in a previous report (22). Resection was conducted using 3-day-old seedlings grown on 1/2 MS medium. Seedlings were placed on 1/2 MS medium with 5× agar and loaded onto a dissecting microscope stage for root tip removal (22). According to the excision position, the resection method was classified into 4 types, namely, the I, II, III and IV types, involving the removal of the QC, 1/2 the meristem, 3/4 the meristem and the whole meristem, respectively. After resection, seedlings were quickly stained with 10 μg/μl propidium iodide (PI) to determine the resection type under a confocal microscope. Finally, the seedlings were moved onto 1/2 MS medium containing 50 μM ampicillin for antibiosis.

To apply the RALF33 peptide to the resected stumps, RALF33 diluted in 1/2 MS liquid medium was dropped onto a small filter paper piece. The resected roots were covered with paper pieces containing 200 nM RALF33, and the dishes were returned to the growth chamber for 72 h. The seedlings were then ready for examination of regeneration.

## ACKNOWLEDGEMENTS

This work was supported by grants from the National Natural Science Foundation of China (NSFC-32070769, 31871396), supported by the science and technology innovation Program of Hunan Province (No.2020WK2014, 2022WK2007, 2021JJ10015), and the China Tobacco Hunan Industrial Co.,Ltd. Research Project (KY2021YC0001).

## AUTHOR CONTRIBUTIONS

F. Yu., W.K. Zhou., and Q.J. Xie conceived the project; Q.J. Xie, W.K. Zhou and F.Yu. designed research; Q.J. Xie, W.J. Chen, S.L. Ouyang and X.N. Wang performed research; F. Xu, Y.R. Wang and L.F. Mao analyzed data; Q.J. Xie and F.Yu. wrote the paper; all authors reviewed and approved the manuscript for publication.

## DECLARATION OF INTERESTS

The authors declare no competing interests.

## DATA AVAILABILITY

The data that support the findings of this study are available from the corresponding author upon reasonable request.

## Notes

### Competing Interest Statement

The authors have declared no competing interest.

