## Supplementary material for "Wounding promotes root regeneration through a cell wall integrity sensor, the receptor kinase FERONIA": SI

**Supporting Information Text**

### Supplementary Information Materials and Methods

**Confocal microscopy and fluorescence quantification**

In this study, we used a Carl Zeiss LSM880 confocal microscope. Unless otherwise specified, the imaging parameters for different trials or time points were identical. We used a 488 nm wavelength argon laser to excite GFP and a 541 nm wavelength to excite PI. When GFP and PI were scanned in parallel, the detection spectrum was set from 490 to 540 nm to avoid signal disturbance from PI. Raw data for fluorescent images were manipulated using Fiji software, wherein membranes were outlined automatically with the same threshold gray value. Z stack images were taken with a step interval of 0.5 μm and reconstructed by Fiji software.

**5-ethynyl-2'-deoxyuridine (Edu) staining**

Five-day-old WT and *fer-4* seedlings grown forehead on 1/2 MS medium were transferred into 1/2 MS liquid medium that containing 10 μM Edu. Seedlings were then incubated in dark in the same growth chamber for 30 min. After incubation, seedlings were fixed with 4% paraformaldehyde for 30 min. The fixation was terminated via adding 50 μl 2 mg/ml glycine. Follow the manufacture of the EdU Staining Proliferation Kit (iFluor 488) (ab219801) for subsequent Click-iT reaction and imaging processes.

**GUS staining**

The GUS staining was performed as described in a previous publication (1).

**Western blotting and antibody preparation**

Western blotting (WB) was performed to assess protein levels in this research. For this purpose, at least 30 μl of seeds were sown at high density on 1/2 MS medium for convenience of treatments or sampling. The method for subsequent experiments were modified according to the specific goals of the experiments.

To examine the effect of RALF33 on TPL/TPR abundance, 5-day-old Col-0 and *fer-4* seedlings were bundled, and the roots were soaked in 1/2 MS liquid medium containing 200 nM RALF33. After incubation for a given duration, the shoots were removed, and the roots were minced for the WB assay. If CHX was needed, the samples were treated with 100 μM CHX (alone or together with RALF33) before sampling.

A 365-amino acid fragment from the N-terminus of TPR1 (2) was used as an antigen to generate the polyclonal anti-TPL/TPR antibody.

**Single-cell RNA-seq**

The raw data and detailed data-processing workflow for single-cell RNA-seq were described in a previous publication (Xu et al., 2022, in preparation).

**RNA extraction and qRT–PCR**

Total RNA was extracted following the manufacturer's protocol for a total RNA isolation kit (RNeasy, Qiagen, No. 74904). The isolated RNA was used to synthesize the first-strand cDNA using ReverTra Ace qPCR RT Master Mix (with gDNA remover; TOYOBO, No. FSQ-301). qRT–PCR was conducted using Realtime PCR Master Mix (TOYOBO, No. QPK-101). Each sample had 3 biological replicates. We used *ACTIN2* as a reference to normalize gene expression levels. The relative expression level compared with that of *ACTIN2* was calculated using the $2^{-\Delta\Delta CT}$ method (3).

**Yeast two-hybrid assay**

The Y2H assay procedure was based on a previous publication (4). In brief, the coding sequence of TPL/TPRs was cloned into the pGADT7 vector for fusion with the AD. The cytosolic domain of FER was cloned into pGBKT7, which contained a BD. TPL/TPR-AD (e.g., TPR1-AD) was cotransformed with FER-CD-BD into yeast AH109 cells. The transformants were diluted with deionized water and dropped onto synthetically defined (SD) medium lacking leucine (SD/-Leu) with an additional 20 mM 3-amino-1,2,4-triazole to evaluate the interaction.

**Split-luciferase complementation assay and subcellular location detection using *N. benthamiana*.**

The split-luciferase complementation assay was based on a previous publication with slight modification (5). Briefly, the *A. tumefaciens* strain GV3101 harboring the pCambia1300-FER-CD-cLUC and pCambia1300-cLuc-TPR1 plasmids was coinfiltrated into *N. benthamiana* leaves and incubated for 2 days in the dark. A 1 mM luciferin solution was smeared onto the infiltrated site on leaves for imaging with a VILBER Newton 7.0 *in vivo* imaging system. To detect the subcellular location of TPR1, full-length CDS of TPR1 was cloned into the pDT7 plasmid, thus to fuse the TPR1 with a GFP tag. The *A. tumefaciens* strain GV3101 harboring the pDT7-TPR1-GFP was coinfiltrated into *N. benthamiana* leaves, and followed by same procedure of incubation of imaging that described in precedent paragraph.

**GST pull-down assay and in vitro phosphorylation**

The C-terminus of FER (225-808 aa) was cloned into pGEXT-4T-1, which harbors a GST tag at the C-terminus, to generate FER-CD-GST. TPR1-Myc was immunoprecipitated from TPR-Myc seedlings using anti-Myc magnetic beads (Bimake, No. B 26302). In detail, 10 μl of serum (containing TPL/TPR antibody) was preincubated with protein A/G at 4°C for 4 h. The beads were washed 3 times with PBST (1×PBS with 0.1% Tween) to remove the excess serum. Then, 0.1 g of 7-day-old TPR1-Myc seedlings was ground to a fine powder, and 1 ml of NEB-T lysis buffer (20 mM HEPES [pH 7.5], 40 mM KCl, 1 mM EDTA, 1 mM PMSF, 1% protease inhibitor cocktail, and 1% Triton X-100) was added to extract proteins at 4°C. The extracts were then centrifuged twice at 16 000 × g at 4°C for 10 min. The supernatant was aspirated and transferred to the prepared beads. The mixture was incubated overnight and washed 3 times with NEB-T wash buffer (with a modified Triton X-100 concentration of 0.2%). The protein was eluted with 0.1 M glycine HCl (pH 3). For the pull-down assay, TPR1-Myc was incubated with GST beads coupled with FER-CD-GST in TBS (50 mM Tris-HCl [pH 7.5], 150 mM NaCl). The beads were then washed 3 times with TBS and boiled with SDS loading buffer, followed by a standard WB assay.

For the *in vitro* phosphorylation assay, the N-terminus and C-terminus of TPR1 with a 6×His tag were expressed with the backbone of pET32a in *E. coli* BL21 cells (0.5 mM IPTG, 110 rpm, 16°C overnight). By following a previously published protein purification procedure (4), we obtained the TPR1-N-His and TPR1-C-His proteins. The kinase domain of FER was generated as previously described (6). In vitro phosphorylation assays were performed according to the protocol described in a previous report (6).

**Co-IP assay**

A total of 0.1 g of 5-day-old FER-Flag transgenic seedlings (6) were sampled, ground, extracted and immunoprecipitated via a specific step in the above passage (IP of TPR1-Myc) using anti-Flag (Bimake, No. B26102) magnetic beads. The beads were washed and boiled for WB analysis. The TPL/TPR antibody was used to detect the interaction.

**Chip-qPCR analysis**

ChIP-qPCR was conducted according to the description in a previous report with modifications (6). In brief, two grams of 5-day-old Col-0 seedlings were cross-linked in 20 mL of 1.0% formaldehyde under vacuum twice for 8 min. After crosslinking, 2 M glycine was added to quench the crosslinking at a final concentration of 0.125 M. After cleaning and nuclear extraction, the precipitate was resuspended and sonicated (80 watts, 5 min). TPL/TPRs were immunoprecipitated using the anti-TPL/TPR antibody (see descriptions in “Western blotting and antibody preparation”) that was prebound to the protein A/G magnetic beads (Bimake, No. B23202), and mouse serum was used as an IP control. After overnight IP, the beads were washed with low-salt, high-salt, LiCl and TE washing buffers (7), and the crosslinking was subsequently revised by adding 20 μl of 5 M NaCl and incubating at 65°C overnight. After decrosslinking, protease-K was used to remove proteins. DNA was extracted by using phenol/chloroform/isoamyl alcohol and precipitated by using excessive 100% ethanol at -80°C for 4 h. Finally, the DNA was dissolved in 20 μl of $ddH_2O$, and the fold enrichment of the given chromatin fragments was determined. The promoter of *ERF115* was detected from -2280 to -2157 bp from the transcription start site.

**Statistical analysis**

Three biological replicates with at least three technical replicates were used in each assay. The mean and standard deviation (SD) values of the replicates were calculated using Microsoft Excel™ software (Microsoft Corp., Redmond, WA, USA). One-way analysis of variance (ANOVA) (Tukey’s HSD test) and Student’s t test were conducted using SPSS software (ver. 23.0; IBM China Company Ltd., Beijing, China) to determine the statistical significance. $p < 0.05$ was considered to indicate a significant difference.

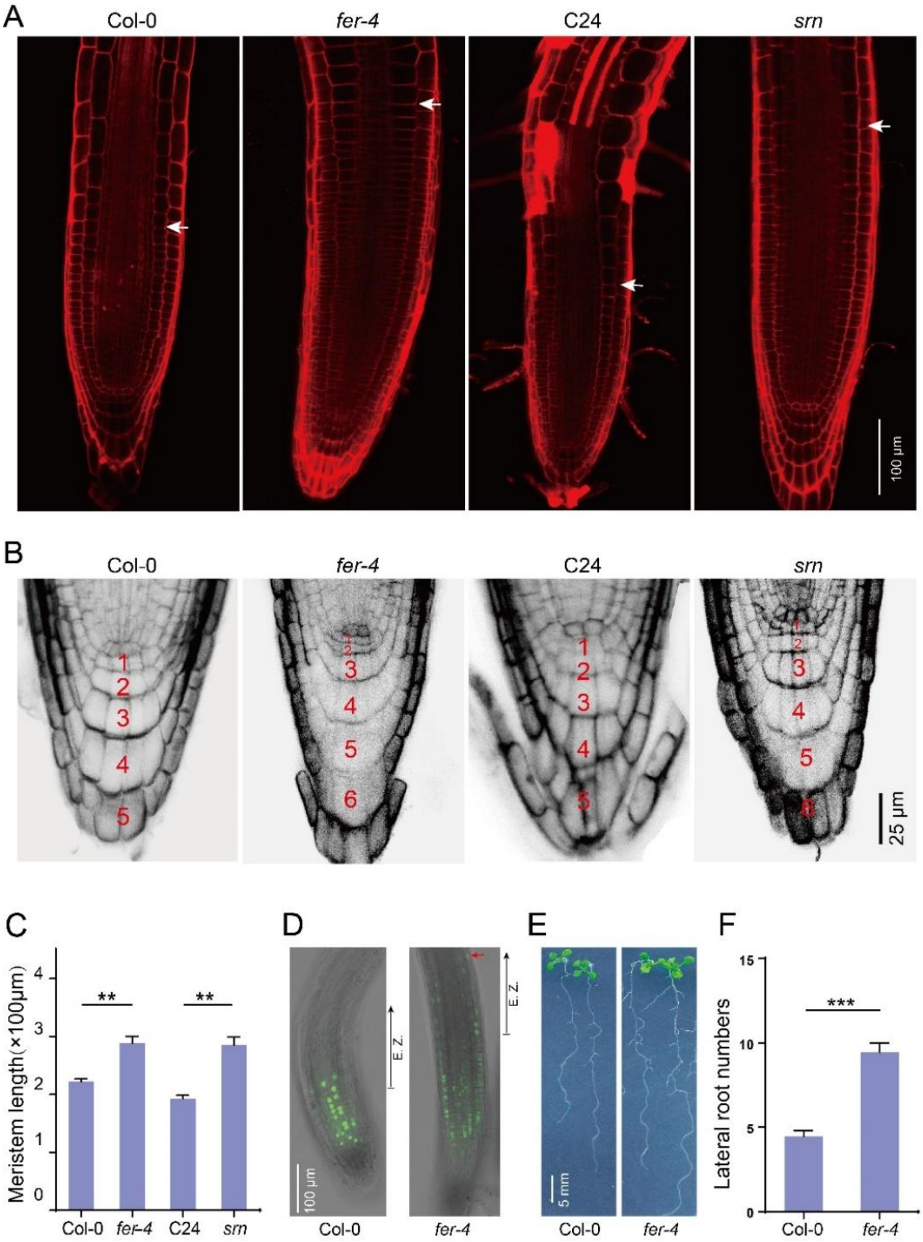


**Fig. S1 regeneration-related phenotypes controlled by *FER*.**

(A) Representative confocal images showing the meristems of Col-0, *fer-4*, C24 and *srn*. The ends of the meristems are indicated by white arrowheads.

(B) Representative images presenting the columella cells of Col-0, *fer-4*, C24 and *srn*. Red numbers indicate the layers of columella cells (including columella stem cells).

(C) Quantification of meristem length of Col-0, *fer-4*, C24 and *srn.* Bars represent the mean ± SE of at least 10 technical replicates per genotype (Student's t test, **$p < 0.01$).

(D) EDU staining of Col-0 and *fer-4*. E. Z. refers to the elongation zone.

(E) Representative images presenting the phenotypes of the lateral roots of Col-0, *fer-4*, C24 and *srn*.

(F) Lateral root numbers of Col-0, *fer-4*, C24 and *srn.* Bars represent the mean ± SE of at least 15 independent seedlings per genotype (Student's t test, ***$p < 0.001$).

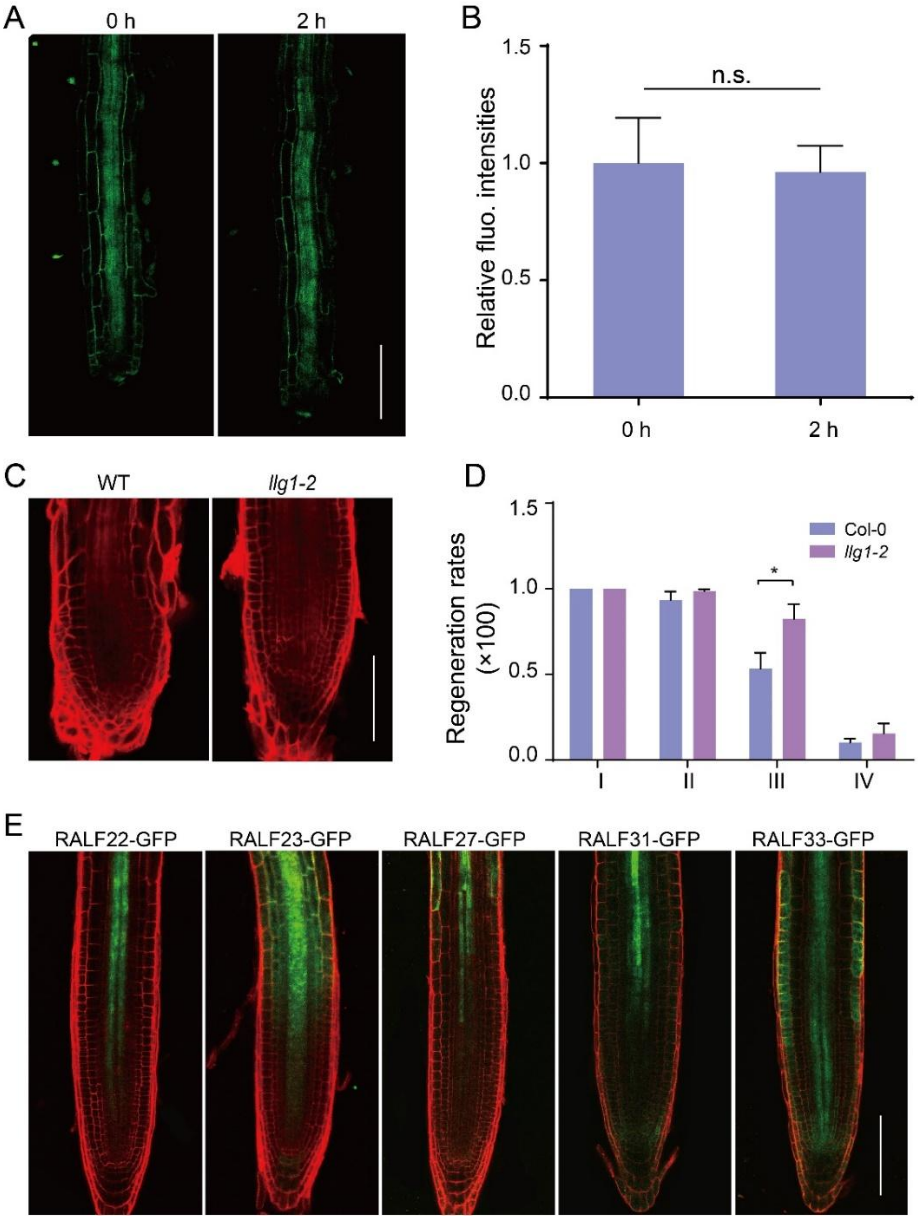


**Figure S2: FER-regulated regeneration requires its binding to ligand.**

(A) Confocal images of FER-GFP after resection. Type III resection was conducted here.

(B) Relative fluorescence intensity of FER-GFP seedlings described in (A). Bars represent the mean ± SE of 3 independent experiments, with at least 15 technical replicates for each trial (Student's t test, n.s., not significant).

(C) Representative roots of the WT and *llg1-2* 3 days after type III resection.

(D) Regeneration frequency of roots described in (C). Bars represent the mean ± SE of 3 independent experiments, with at least 15 technical replicates for each trial (Student's t test, *$p < 0.05$).

(E) Expression pattern of meristem-expressed *RALFs*. The images presented here represent PI-stained *pRALF::RALF-GFP* reporter lines. For clear visualization, the imaging settings were not necessarily identical. Bar = 100 μm.

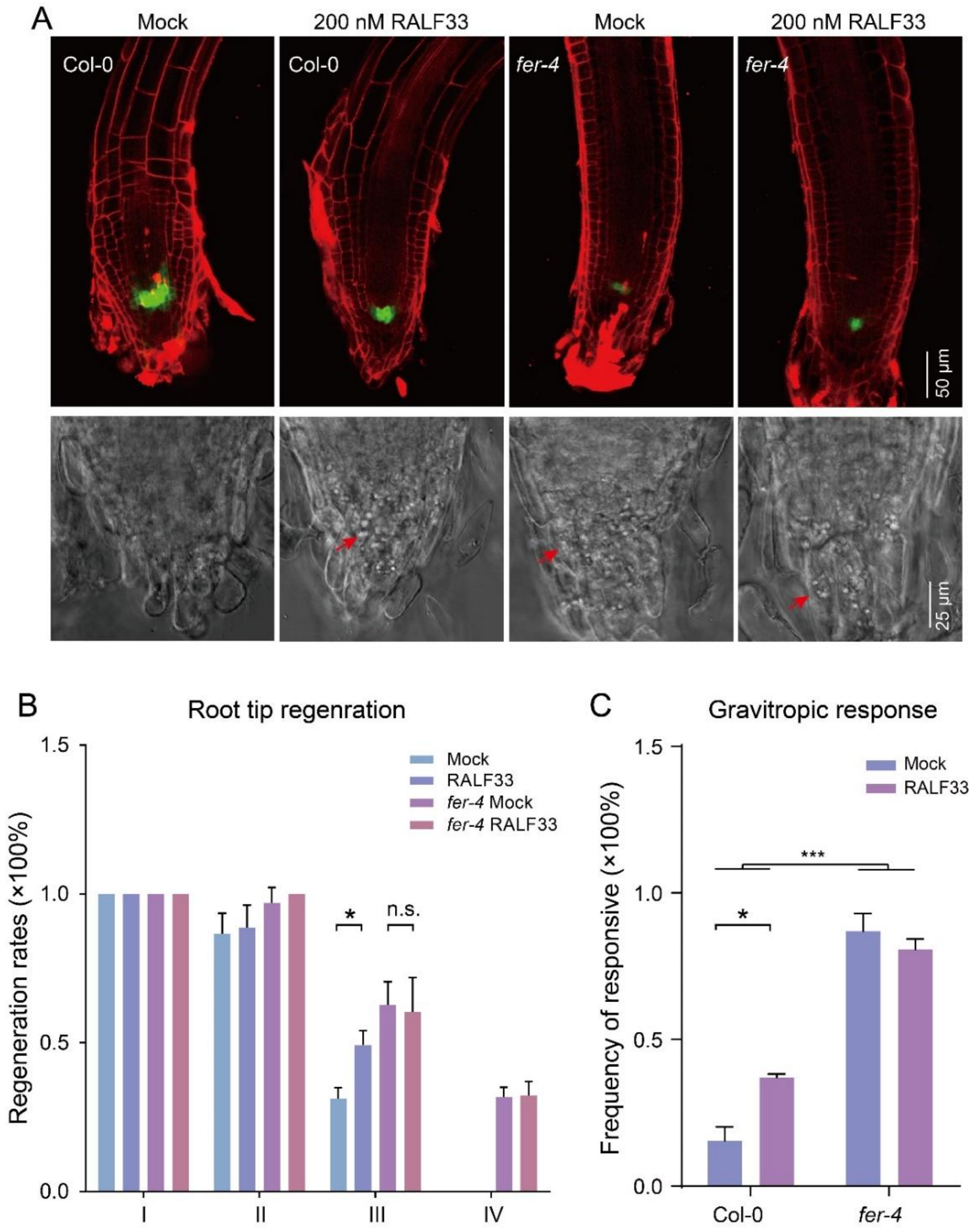


**Figure S3: RALF33 stimulates regeneration through FER.**

(A) Confocal images of WOX5::GFP/Col-0 and WOX5::GFP/*fer-4* treated with/without 200 nM RALF33 after resection. Type III resection images are shown here. Green signals indicate *WOX5-GFP*. The lower panels of the bright field images show the columella containing starch granules (indicated by red arrows).

(B) Regeneration rate of Col-0 and *fer-4* treated with/without 200 nM RALF33. Bars represent the mean ± SE of 3 independent experiments, with at least 15 technical replicates per trial (Student's t test, n.s., not significant; *$p < 0.05$).

(C) Frequency of gravitropism-responsive seedlings of Col-0 and *fer-4* after type III resection with/without 200 nM RALF33. Bars represent the mean ± SE of 3 independent experiments, with at least 15 technical replicates per trial (Student's t test, n.s., not significant; *$p < 0.05$ ;***$p < 0.001$).

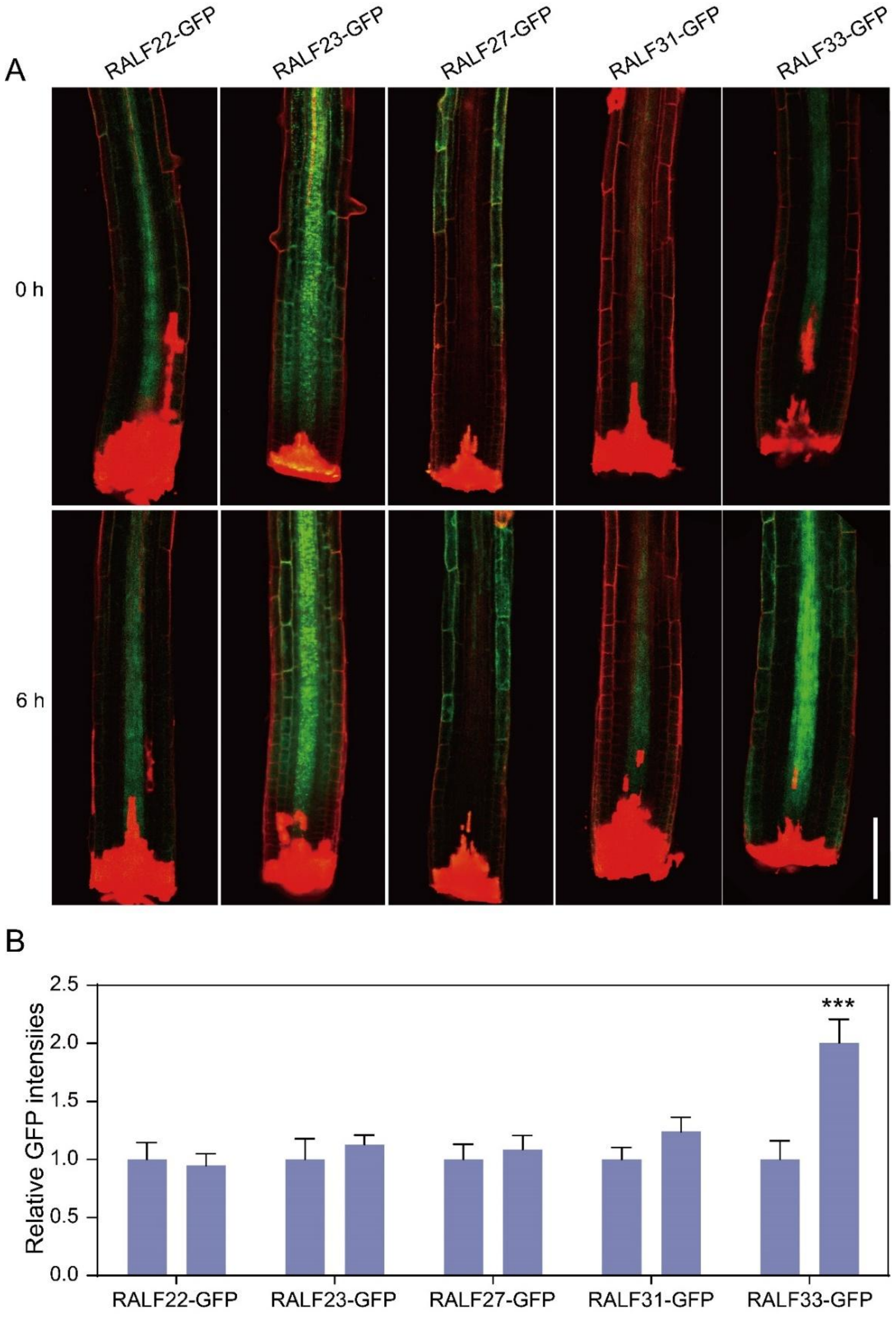


**Figure S4: Dynamics of RALF abundance in response to wounding.**

(A) Confocal images of RALFs-GFP 6 h after type III resection.

(B) Relative fluorescence intensity of RALF-GFP seedlings described in (A). The data are presented as the mean ± SD, and each trial had at least 15 replicates (Student's t test, ***$p < 0.001$).

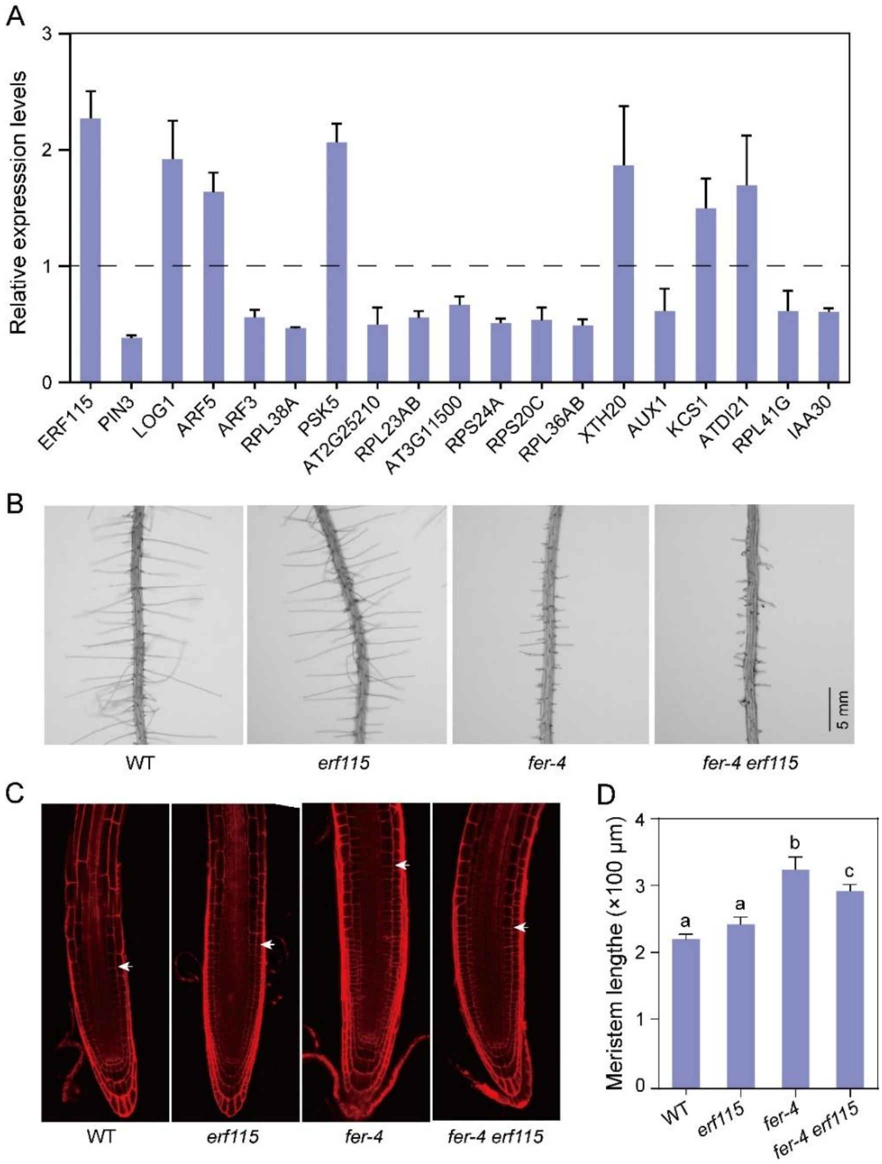


**Figure S5: ERF115 is a potential downstream factor of FER that regulates regeneration.**

(A) Relative expression levels of regeneration-related DEGs in the low-differentiation cells in the stele of *fer-4*. The expression levels were normalized to that in Col-0. The black dashed line indicates normalized expression levels in Col-0. Expression levels were obtained from scRNA-seq data of Col-0 and *fer-4* roots, with 3 biological replicates. The expression level in *fer-4* was normalized to that in Col-0.

(B) Representative root hairs of Col-0, *fer-4*, *erf115*, and *fer-4 erf115*. Seedlings presented here were 5 days old. Roots grown on 1/2 growth medium were imaged on a stereomicroscope.

(C) Confocal images of the root tips of 5-day-old Col-0, *fer-4*, *erf115*, and *fer-4 erf115*. The white arrow indicates the boundaries of the meristem.

(D) Meristem lengths of 5-day-old Col-0, *fer-4*, *erf115*, and *fer-4 erf115*. The data shown indicate the mean ± SE of at least 10 technical replicates per genotype. The different lowercase letters indicate statistical significance (one-way ANOVA).

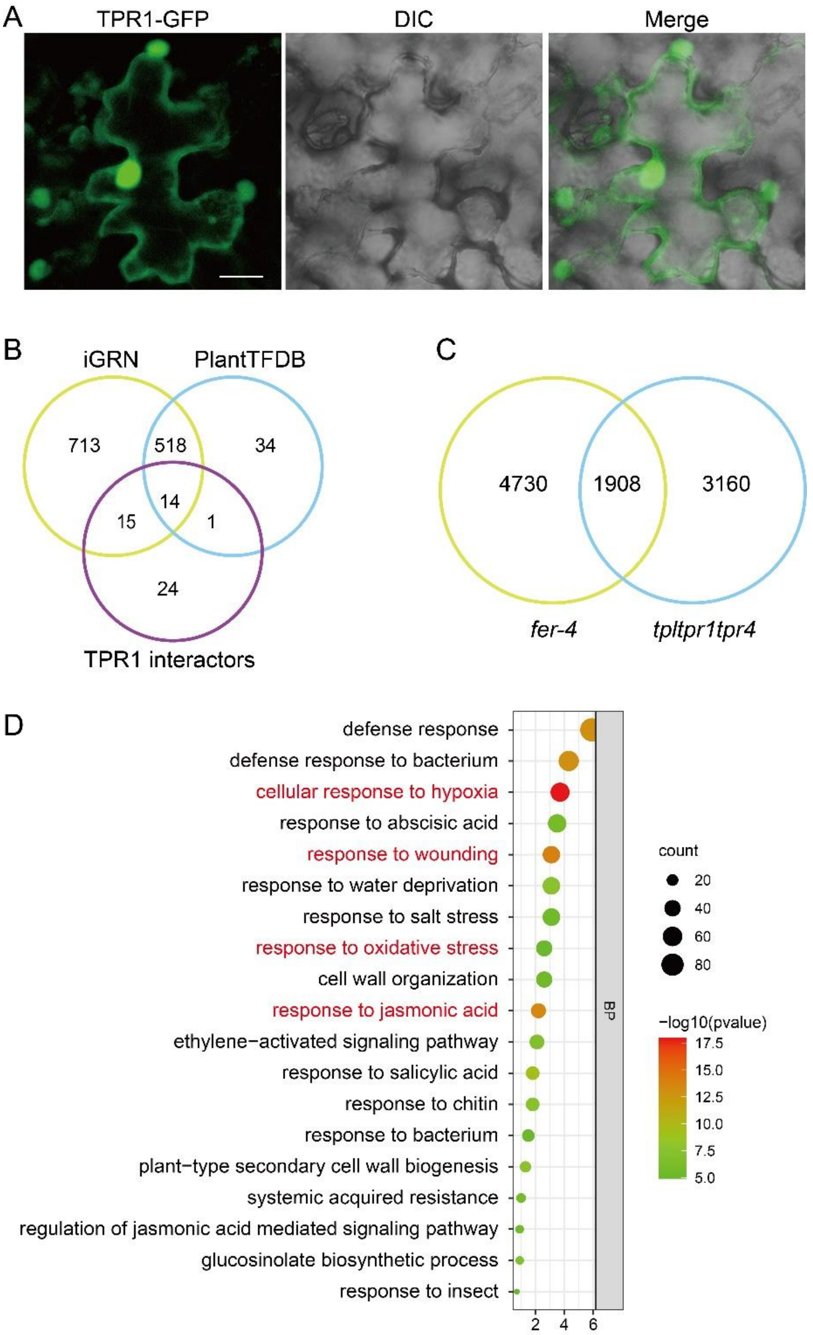


**Figure S6: Subcellular location of TPR1, and bioinformatic analysis of the possibility that TPL/TPRs function downstream of FER.**

(A) Subcellular location of TPR1. Fusion protein of GFP-tagged TPR1 carried by pDT7 plasmid was expressed in *N. benthamiana* leaves, and detected using confocal microscope. Bar = 20 µm.

(B) Venn diagram of enriched TFs of regeneration-related genes and TPR1 interactors. iGRN and PlantTFDB (See hyperlink in main text) represent two independent databases for TFs enrichment analysis.

(C) Venn diagram of DEGs of *fer-4* v.s. Col-0 and *tpltpr1tpr4* v.s. Col-0.

(D) GO enrichment of the overlapping genes described in (B).

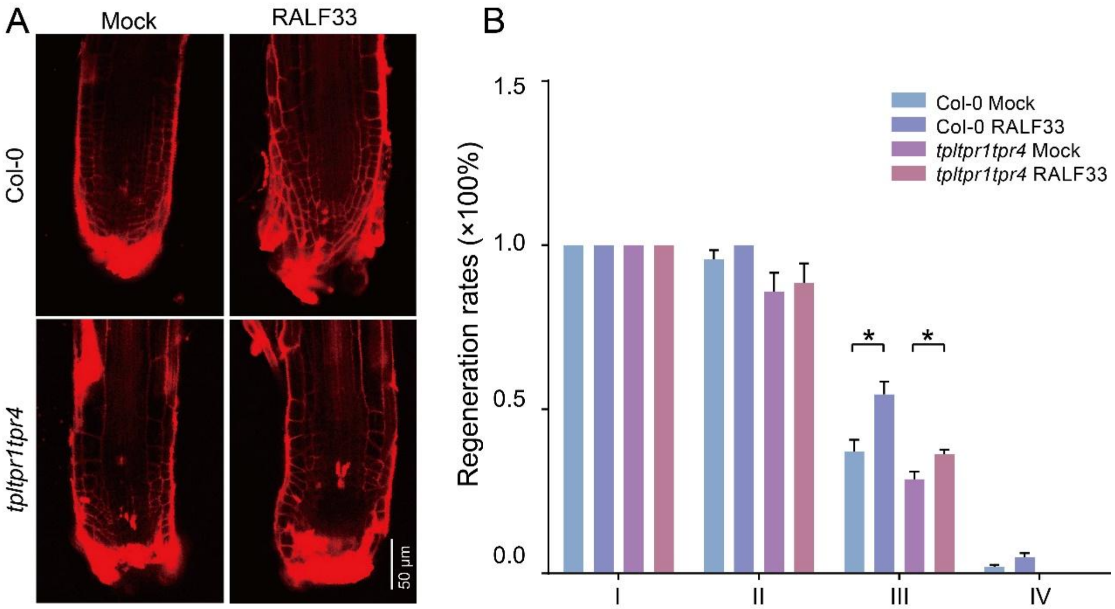


**Fig S7 Characterization of the genetic relationship of RALF33 and TPL/TPRs.**

(A) Col-0, *tpltpr1tpr4* and *TPR1-OE* seedlings following RALF33 treatment after resection. 200 nM RALF33 were added after type III resection, the roots were stained and imaged 72 h later.

(B) The regeneration rates of Col-0 and *tpltpr1tpr4* following RALF33 treatments. Bars represent the mean of three replicated trials ± SE. Each trial is with at least 15 independent seedlings (Student's t test, *$p < 0.05$).
